# A Minimal Medium for Culturing Maize Root-Associated Microbes Based on a Plant Growth Medium

**DOI:** 10.1101/2025.07.07.663485

**Authors:** Anna-Katharina Garrell, Simina Vintila, Henry Phillip, Ashley E. Beck, Manuel Kleiner

**Author notes:** Corresponding author: Manuel Kleiner.

## Abstract

Plant-associated microbiota play a critical role in host resilience to both abiotic and biotic stresses. However, understanding the underlying mechanisms of interaction within these communities, as well as between these communities and their hosts, remains challenging due to the complexity and dynamic nature of plant-associated microbiota. Synthetic microbial communities (SynComs) can serve as experimentally tractable models to establish a fundamental understanding of plant-microbiota interactions. Here, we report the development of a defined, minimal growth medium for a well-characterized seven-member maize root SynCom. The medium is based on a standard plant-growth medium to enable its use for both *in vitro* and *in planta* studies. Using genome-scale metabolic modeling and auxotrophy prediction, we identified key nutrient requirements of each of the seven species and optimized the medium composition to support bacterial growth. This minimal medium enables controlled investigation of microbial physiology, metabolite exchange, and community interactions, and lays the foundation for scalable *in vitro* and *in planta* experiments, facilitating future research on plant microbiome functions.

## Introduction

Plant-associated microbiota can provide a wide range of benefits to their plant hosts, including enhancing resilience to abiotic stresses (Doty 2011; Li et al. 2006; Poudel et al. 2021; Yang et al. 2009) - such as nutrient deficiency and drought - and biotic stresses, such as pathogen pressure (Bernal et al. 2017; Elsayed et al. 2020; Haas and Défago 2005). Given these roles, microbial communities are increasingly recognized as a promising component of sustainable agricultural strategies. Plant microbiota are, however, highly diverse and are influenced by a wide range of factors including host genotype and tissue, soil type, agricultural management practices, drought and temperature conditions, and interactions with other organisms (Coleman-Derr et al. 2016; Fitzpatrick et al. 2018; Mo et al. 2024; Wipf et al. 2021). Due to this variability, as well as their complex and dynamic natures, the multi-level interactions in these systems (between members of the microbiota and between the microbiota and their hosts) remain poorly understood. Recent studies have used various -omics techniques, transposon mutagenesis, and metabolic modeling to begin untangling these interactions to understand the underlying mechanisms (Cole et al. 2017; Hemmerle et al. 2022; Lidbury et al. 2022; Schäfer et al. 2023; Vannier et al. 2023). In this work, synthetic communities (SynComs) have proven particularly useful as tractable models for investigating plant-microbiota interactions (Marín et al. 2021; Vorholt et al. 2017).

One such SynCom was published in 2017 for maize by Niu *et al*. (2017) and consists of seven bacterial species: *Stenotrophomonas maltophilia* (SMA), *Brucella pituitosa* (BPI), *Curtobacterium pusillum* (CPU), *Enterobacter ludwigii* (ELU), *Chryseobacterium indologenes* (CIN), *Herbaspirillum robiniae* (HRO), and *Pseudomonas putida* (PPU). These seven species consistently colonized maize roots up to 15 days and were shown to inhibit the fungal pathogen *Fusarium verticillioides in planta*. Additionally, ELU was hypothesized to be a keystone species, as its removal resulted in a collapse of the community structure. This SynCom has since been used to investigate microbe-dependent heterosis (Wagner et al. 2021) and metabolism in a community setting (Krumbach et al. 2021), prompting the development of additional tools and methods, including protein extraction methods and a modular cloning toolkit (Parnell et al. 2023; Salvato et al. 2022; van Schaik et al. 2023). Additionally, the seven species are available from the public microbial strain repository DSMZ enabling the broader community to use this SynCom (Garrell et al. 2025). As such, this community is proving to be a valuable system to investigate a variety of questions, including colonization mechanisms, community assembly, microbe-microbe interactions, and pathogen resistance.

In this study, we used genome-scale metabolic models to guide the development of a defined, minimal medium that supports the growth of each of the maize SynCom members individually. Having a defined minimal medium will allow for precise manipulation of nutrients, enabling investigation into microbial metabolism and physiology, metabolite exchange, resource competition, and other interactions within the community. A minimal medium will also provide a basis for comparing microbial functions between free-living and plant-associated growth.

## Results

To develop this medium, we established four constraints: 1) The medium must be defined so that individual components can be manipulated to study the impacts of specific compounds on bacterial growth, functions, and interactions. 2) The medium must be minimal, so as to reduce complexity and minimize confounding results in any future experiments. 3) The medium should be as similar as possible to a plant growth medium to enable more direct comparisons between *in vitro* and *in planta* growth. 4) The medium must support growth of each of the seven species in monoculture, so that hypotheses may be tested on individual species as well as various combinations of SynCom members.

Based on the criteria outlined above, we first aimed to establish the base medium and primary carbon sources by investigating bacterial growth in two plant growth media - Murashige-Skoog (MS) or Hoagland’s - with glucose and malate (Fig. 1), as well as sucrose (not shown). MS and Hoagland’s were selected as they have been frequently used for maize growth (Kaya et al. 2013; Niu et al. 2017; Wagner et al. 2021; Zhao et al. 2003), satisfying the requirement that the medium be as similar as possible to the inoculation medium used in experiments with plants. The carbon sources were selected due to their presence in plant root exudates (Carvalhais et al. 2011; Farrar et al. 2003; Jones 1998; Zhalnina et al. 2018). For this initial screening experiment to determine base medium and carbon sources, we used two replicates.

**Figure 1.**
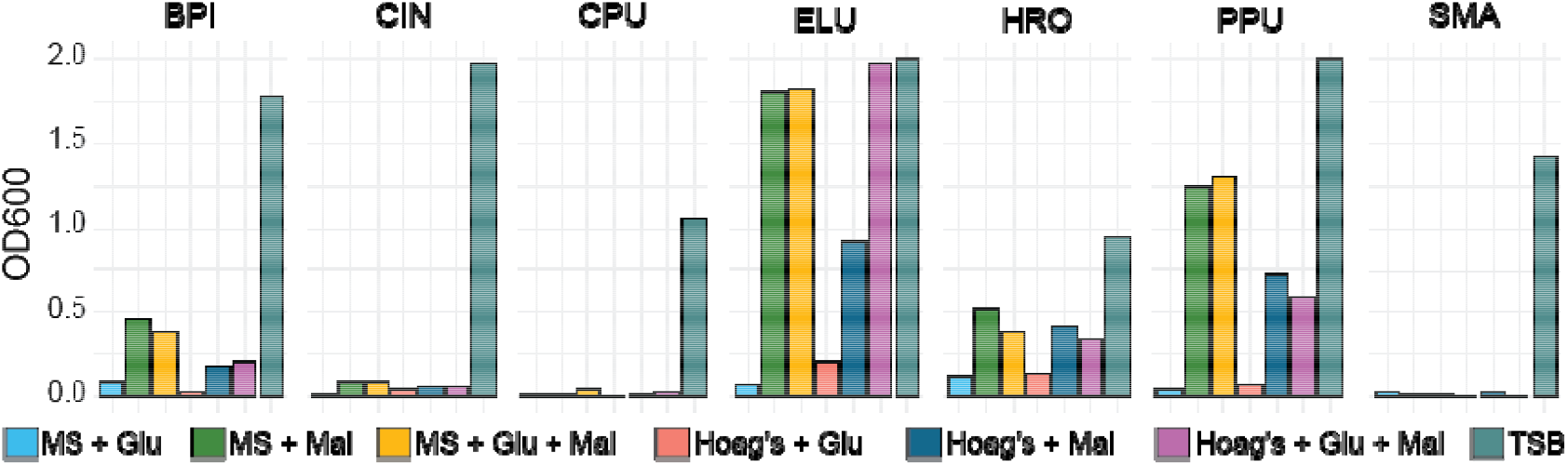
Growth of SynCom members after 24 hours in 0.5x Murashige-Skoog (MS) and Hoagland’s with glucose and/or malate (n = 2). “MS + Glu” - 0.5x MS with glucose; “MS + Mal” - 0.5x MS with malate; “MS + Glu + Mal” - 0.5x MS with glucose and malate; “Hoag’s + Glu” - Hoagland’s with glucose; “Hoag’s + Mal” - Hoagland’s with malate; “Hoag’s + Glu + Mal” - Hoagland’s with glucose and malate; “TSB” - tryptic soy broth.

We found that BPI, ELU, HRO, and PPU grew best in media augmented with malate irrespective of glucose availability (Fig. 1), particularly when compared to their growth in the complex medium TSB. BPI and PPU showed slightly higher growth in MS than in Hoagland’s, while HRO and ELU did not show a strong preference for one base medium over the other. CIN, CPU, and SMA did not grow well in either of the two initial media, though CPU showed a very slight OD increase in 0.5x MS with glucose and malate, and SMA showed a very slight OD increase in 0.5x MS with glucose and Hoagland’s with malate. None of the seven species grew well in media augmented with sucrose.

Given that there was a preference by some species for MS over Hoagland’s, a strong preference for malate by some species, and a slight preference for glucose by some species, we decided to proceed with MS with glucose and malate for further optimization. We used the “Predict Genome Auxotrophies” tool in KBase to predict amino acid and vitamin auxotrophies for each species to guide further nutrient supplementation (Table 1). We then augmented the MS-based medium with amino acids and vitamins for which at least one species was auxotrophic to determine whether their addition would increase bacterial growth.

**Table 1.** Amino acid and vitamin auxotrophies predicted for seven bacterial species by the “Predict Genome Auxotrophies” tool in KBase. “+” indicates a predicted auxotrophy.

|  | BPI | CIN | CPU | ELU | HRO | PPU | SMA |
| --- | --- | --- | --- | --- | --- | --- | --- |
| Arg | + | + | + |  | + |  |  |
| Pro | + | + |  |  | + |  |  |
| Ser |  |  |  | + |  |  |  |
| Asn |  |  |  |  | + | + | + |
| Ile | + |  | + |  | + | + | + |
| Leu | + |  | + |  | + | + | + |
| Lys | + | + | + |  | + | + |  |
| Cys |  | + | + | + |  |  | + |
| Gln |  |  |  |  |  | + |  |
| Niacin (B3) |  |  |  |  |  |  | + |
| Cobalamin (B12) |  |  | + |  |  |  |  |
| Riboflavin (B2) |  |  |  | + |  |  |  |

As a result, we found that the addition of Arg, Asn, Cys, Gln, Ile, Leu, Lys, Pro, Ser, niacin (B3), cobalamin (B12), and riboflavin (B2) resulted in a significant increase in BPI, CIN, CPU, and SMA growth, and to some extent, in PPU (Fig. 2A), although not significant. However, despite the increase, we still were not able to obtain much growth for CPU, particularly when compared to its growth in TSB.

**Figure 2.**
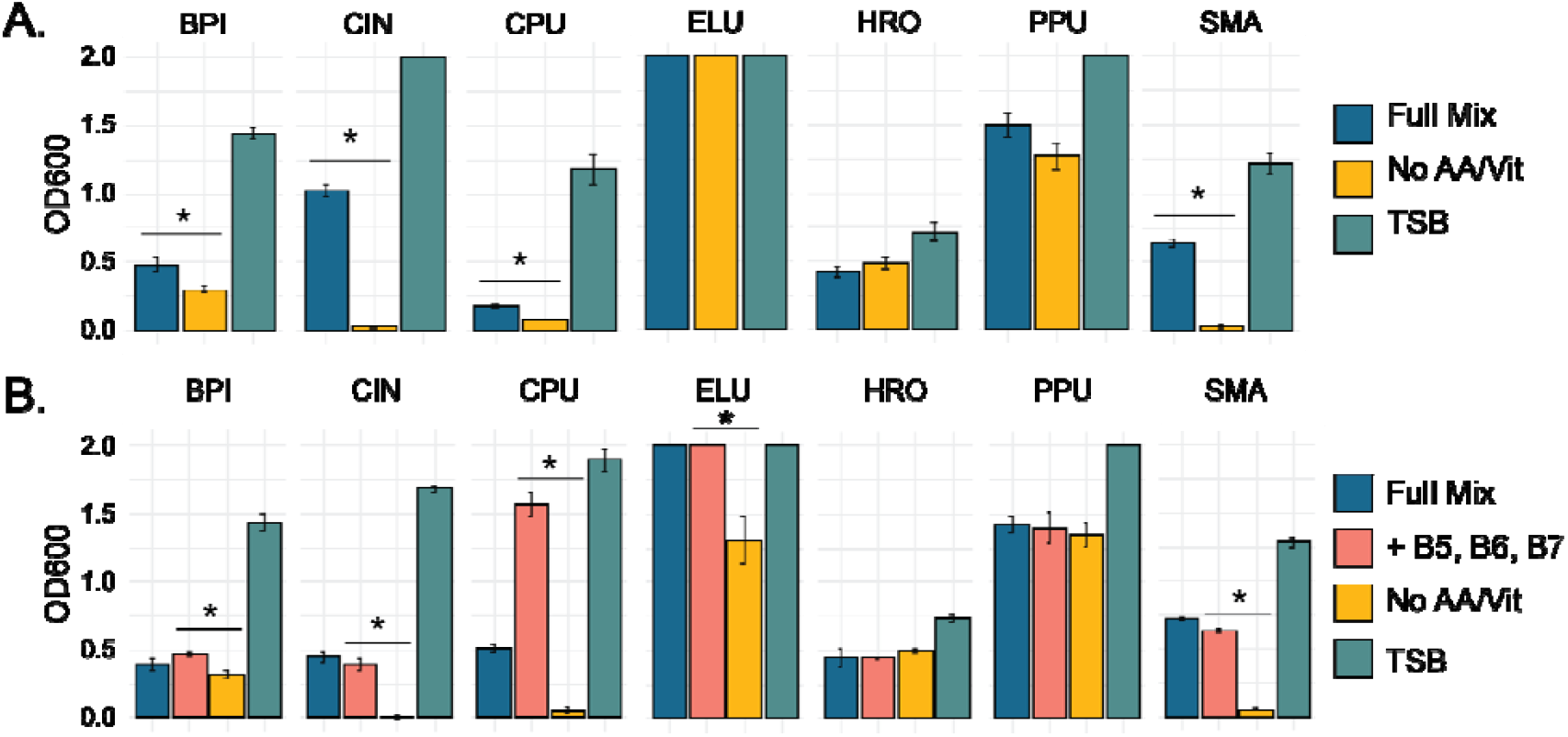
Growth of SynCom species in 0.5x MS, glucose, and malate augmented with additional amino acids and vitamins (n = 3). A) OD600 measured after 24 hours of growth of bacterial species when grown in three different media. “Full Mix” refers to the minimal medium augmented with Arg, Pro, Ser, Asn, Ile, Leu, Lys, Cys, Gln, niacin (B3), cobalamin (B12), riboflavin (B2). “No AA/Vit” refers to the minimal medium containing no vitamins or amino acids. “TSB” refers to tryptic soy broth. (n = 3). Asterisk indicates p < 0.05 (Student’s T-Test) when growth in the “Full Mix” was compared to growth in the “No AA/Vit” medium. Error bars indicate standard deviation. B) OD600 of bacterial species when grown in four different media (separate experiment from panel A). “Full Mix”, “No AA/Vit”, and “TSB” are the same media as described in (A). “+ B5, B6, and B7” refers to the Full Mix medium with the addition of pantothenate (B5), pyridoxine (B6), and biotin (B7). Asterisks indicate p < 0.05 (Student’s T-Test) when growth in the “+ B5, B6, and B7” was compared to growth in the “No AA/Vit” medium. Error bars represent standard deviation.

Taking another approach to identify potential auxotrophies in CPU, we constructed a metabolic model for CPU in KBase and performed gapfilling on the minimal medium (MS with glucose and malate), either with or without the amino acids and vitamins listed in Table 1. This step identified gaps in essential pathways and added any reactions that were missing from the genome to permit growth on the specified medium. We then examined the gapfilled reactions to determine biosynthetic pathways that had not previously been identified by the “Predict Genome Auxotrophies” tool. We found that CPU did not encode all genes necessary for biotin (B7), pyridoxine (B6), and pantothenate (B5) biosynthesis or glycine metabolism. We therefore started by adding the three vitamins to the medium and observed a significant improvement in CPU growth (Fig. 2B). We did not test the effect of additional glycine, as we found sufficient growth with the addition of vitamins B5, B6, and B7.

## Discussion

We have developed a defined, minimal medium (Table 2) that supports the *in vitro* cultivation of all seven bacterial species in the maize root SynCom described by Niu *et al*. (2017) As a result, we present a system that will enable targeted, mechanistic investigations of plant-associated microbes under controlled, reproducible conditions.

**Table 2.** Final composition of the defined, minimal medium.

| Component | Final Concentration |
| --- | --- |
| Murashige-Skoog<br>(without sucrose) | 0.5x |
| Malate | 1% w/v |
| Glucose | 1% w/v |
| Arginine | 50 µg/mL |
| Proline | 50 µg/mL |
| Serine | 50 µg/mL |
| Asparagine | 50 µg/mL |
| Leucine | 50 µg/mL |
| Isoleucine | 50 µg/mL |
| Lysine | 50 µg/mL |
| Cysteine | 50 µg/mL |
| Glutamine | 50 µg/mL |
| Riboflavin (B2) | 0.2 µg/mL |
| Nicotinic Acid (B3) | 1 µg/mL |
| Cyanocobalamin (B12) | 0.02 µg/mL |
| Pyridoxine (B6) | 1 µg/mL |
| Pantothenate (B5) | 1 µg/mL |
| Biotin (B7) | 0.01 µg/mL |

We recognize that this medium has limitations and could benefit from further refinement. We did not perform amino acid or vitamin drop-out experiments to confirm whether each was essential to bacterial growth, so the medium could potentially be reduced further. Additionally, for single-species studies, particularly those involving species with few predicted auxotrophies, e.g. ELU, the medium could be simplified further to reflect only the amino acid and vitamin requirements of that given species. Furthermore, while this medium supports monoculture bacterial growth and is anticipated to also support community growth, future work should evaluate its ability to support co-culture growth to enable experiments studying microbe-microbe interactions.

Notwithstanding, the medium presented here provides a platform for future research into microbial physiology, microbe-host, and microbe-microbe interactions. In particular, given the defined nature of this medium, it is well-suited for probing microbial metabolism and nutrient uptake, particularly under nutrient limited conditions as each component can be easily manipulated. Additionally, because the medium is based on Murashige-Skoog, a medium commonly used in maize growth experiments, it will allow for comparative studies between bacterial growth *in vitro* and *in planta*, supporting investigation of plant-microbe interactions. It can additionally be used to investigate microbe-microbe interactions involving nutrient competition and metabolite exchange in co-culture experiments. Importantly, this medium also provides a basis for validating and refining genome-scale metabolic models, facilitating *in silico* simulations of microbial growth and interactions under experimentally relevant conditions. This dual utility - supporting both experimental and computational approaches - will aid in bridging the gap between modeling and experimental validation in microbial ecology.

Ultimately, this defined, minimal medium will support bottom-up mechanistic studies using both wet lab and computational techniques to gain a more holistic, systems-level understanding of plant-associated microbes and microbiota.

## Materials and Methods

### Culturing of SynCom Species

The previously published SynCom used in this study contains the isolates *Stenotrophomonas maltophilia* AA1 (SMA; DSM 114483), *Brucella pituitosa* AA2 (BPI; DSM 114565), *Curtobacterium pusillum* AA3 (CPU; DSM 114566), *Enterobacter ludwigii* AA4 (ELU; previously *E. cloacae*; DSM 114484), *Chryseobacterium indologenes* AA5 (CIN; 114485), *Herbaspirillum robiniae* AA6 (HRO; DSM 114508), and *Pseudomonas putida* AA7 (PPU; DSM 114486). We cultured all isolates as described by Salvato *et al*. (2022) (Salvato et al. 2022). Briefly, we streaked bacterial species from frozen glycerol stocks on selective 0.1x tryptic soy agar (TSA) plates, as described in Niu *et al*. (Niu et al. 2017; Niu and Kolter 2018) and incubated them at 30°C for 48 hours. We inoculated a single colony from each species into 5 mL of tryptic soy broth (TSB) and incubated them in a shaker at 180 rpm at 30°C for 8 hours. 50 µL of this culture was then inoculated into 5 mL of TSB and again incubated at 180 rpm at 30°C for 16 hours. Cells were then washed to remove media components by pelleting 1 mL of the overnight culture at 8000 xg for 8 minutes, resuspended in 1 mL of 1x phosphate buffered saline (PBS; Fisher Scientific BP2944100), and pelleted again. The final pellet was resuspended in 1 mL PBS.

We inoculated 5 mL of the different versions of the minimal media with 50 µL of the washed, resuspended cells (n = 3). We incubated the cultures in a shaker at 180 rpm at 30°C for 24 hours and then measured culture optical density at 600 nm (OD600) using a Biochrom Ultrospec™ 10.

Recipes for defined, minimal media are listed in Supplementary Information. Briefly, stock solutions of vitamins were prepared, filter-sterilized through 0.22 µm filters, and stored in the dark at 4°C. Stock solutions of amino acids were prepared and stored at -20°C. One day before experiments were set up, Murashige-Skoog salts or Hoagland’s (Niu et al. 2016) solutions were added to water and autoclaved at 121°C for 30 minutes. Glucose and malate solutions were prepared separately and filter-sterilized through 0.22 µm filters. A master mix of amino acids was prepared from the stock solutions and filter-sterilized through a 0.22 µm filter. Glucose, malate, vitamins, and amino acids were added to the sterile Murashige-Skoog or Hoagland’s solutions once at room temperature.

### Auxotrophy Prediction and Metabolic Model Generation

Genomes for each SynCom species were re-annotated in KBase (Arkin et al. 2018) with RASTtk (Brettin et al. 2015) using the “Annotate Microbial Genome” tool (Refseq numbers CP018756 (SMA), CP018779-82 (BPI), CP018783-4 (CPU), CP018785 (ELU), CP018786 (CIN), CP018845 (HRO), and CP018846 (PPU)). These annotated genomes were used as input for the “Predict Genome Auxotrophies” tool in KBase to identify expected auxotrophies for each species. KBase narratives are available for each species (at https://narrative.kbase.us/narrative/#, with # ranging from 65186-65192).

A metabolic model was additionally constructed for CPU based on the RASTtk-annotated genome with the “Build Metabolic Model” tool in KBase. Gapfilling was performed simultaneously with model construction using the minimal medium including the suite of experimentally added amino acids (Arg, Asn, Cys, Gln, Ile, Leu, Lys, Pro, Ser) and vitamins (cobalamin, niacin, and riboflavin). The model output was downloaded from KBase as an Excel file to manually identify gapfilled reactions (lacking a genome-annotated reference), indicating additional auxotrophies missed in the original auxotrophy prediction. KBase narratives are available for the model construction and media testing (narrative numbers 92979 and 95197).

## Acknowledgements

We thank all Kleiner lab members for discussions on methods, results, and data analysis. This research was funded by National Institute of Food and Agriculture awards 2021-67013-34537 (AEB and MK) and 2022-67013-36672 (MK), and a Novo Nordisk Foundation award NNF19SA0059360 (MK).

## Literature Cited

1. Arkin, A. P., Cottingham, R. W., Henry, C. S., Harris, N. L., Stevens, R. L., Maslov, S., Dehal, P., Ware, D., Perez, F., Canon, S., Sneddon, M. W., Henderson, M. L., Riehl, W. J., Murphy-Olson, D., Chan, S. Y., Kamimura, R. T., Kumari, S., Drake, M. M., Brettin, T. S., Glass, E. M., Chivian, D., Gunter, D., Weston, D. J., Allen, B. H., Baumohl, J., Best, A. A., Bowen, B., Brenner, S. E., Bun, C. C., Chandonia, J.-M., Chia, J.-M., Colasanti, R., Conrad, N., Davis, J. J., Davison, B. H., DeJongh, M., Devoid, S., Dietrich, E., Dubchak, I., Edirisinghe, J. N., Fang, G., Faria, J. P., Frybarger, P. M., Gerlach, W., Gerstein, M., Greiner, A., Gurtowski, J., Haun, H. L., He, F., Jain, R., Joachimiak, M. P., Keegan, K. P., Kondo, S., Kumar, V., Land, M. L., Meyer, F., Mills, M., Novichkov, P. S., Oh, T., Olsen, G. J., Olson, R., Parrello, B., Pasternak, S., Pearson, E., Poon, S. S., Price, G. A., Ramakrishnan, S., Ranjan, P., Ronald, P. C., Schatz, M. C., Seaver, S. M. D., Shukla, M., Sutormin, R. A., Syed, M. H., Thomason, J., Tintle, N. L., Wang, D., Xia, F., Yoo, H., Yoo, S., and Yu, D. 2018. KBase: The United States Department of Energy Systems Biology Knowledgebase. Nat. Biotechnol. 36:566–569. 10.1038/nbt.4163.

2. Bernal, P., Allsopp, L. P., Filloux, A., and Llamas, M. A. 2017. The Pseudomonas putida T6SS is a plant warden against phytopathogens. ISME J. 11:972–987. 10.1038/ismej.2016.169.

3. Brettin, T., Davis, J. J., Disz, T., Edwards, R. A., Gerdes, S., Olsen, G. J., Olson, R., Overbeek, R., Parrello, B., Pusch, G. D., Shukla, M., Thomason, J. A., Stevens, R., Vonstein, V., Wattam, A. R., and Xia, F. 2015. RASTtk: A modular and extensible implementation of the RAST algorithm for building custom annotation pipelines and annotating batches of genomes. Sci. Rep. 5:8365. 10.1038/srep08365.

4. Carvalhais, L. C., Dennis, P. G., Fedoseyenko, D., Hajirezaei, M.-R., Borriss, R., and von Wirén, N. 2011. Root exudation of sugars, amino acids, and organic acids by maize as affected by nitrogen, phosphorus, potassium, and iron deficiency. J. Plant Nutr. Soil Sci. 174:3–11. 10.1002/jpln.201000085.

5. Cole, B. J., Feltcher, M. E., Waters, R. J., Wetmore, K. M., Mucyn, T. S., Ryan, E. M., Wang, G., Ul-Hasan, S., McDonald, M., Yoshikuni, Y., Malmstrom, R. R., Deutschbauer, A. M., Dangl, J. L., and Visel, A. 2017. Genome-wide identification of bacterial plant colonization genes ed. Xinnian Dong. PLOS Biol. 15:e2002860. 10.1371/journal.pbio.2002860.

6. Coleman-Derr, D., Desgarennes, D., Fonseca-Garcia, C., Gross, S., Clingenpeel, S., Woyke, T., North, G., Visel, A., Partida-Martinez, L. P., and Tringe, S. G. 2016. Plant compartment and biogeography affect microbiome composition in cultivated and native Agave species. New Phytol. 209:798–811. 10.1111/nph.13697.

7. Doty, S. L. 2011. Nitrogen-Fixing Endophytic Bacteria for Improved Plant Growth. In Bacteria in Agrobiology: Plant Growth Responses, ed. Dinesh K. Maheshwari. sBerlin, Heidelberg: Springer, pp. 183–199. 10.1007/978-3-642-20332-9_9.

8. Elsayed, T. R., Jacquiod, S., Nour, E. H., Sørensen, S. J., and Smalla, K. 2020. Biocontrol of Bacterial Wilt Disease Through Complex Interaction Between Tomato Plant, Antagonists, the Indigenous Rhizosphere Microbiota, and Ralstonia solanacearum. Front. Microbiol. 10.

9. Farrar, J., Hawes, M., Jones, D., and Lindow, S. 2003. How Roots Control the Flux of Carbon to the Rhizosphere. Ecology 84:827–837. 10.1890/0012-9658(2003)084[0827:HRCTFO]2.0.CO;2.

10. Fitzpatrick, C. R., Copeland, J., Wang, P. W., Guttman, D. S., Kotanen, P. M., and Johnson, M. T. J. 2018. Assembly and ecological function of the root microbiome across angiosperm plant species. Proc. Natl. Acad. Sci. 115:E1157– E1165. 10.1073/pnas.1717617115.

11. Garrell, A.-K., Cheadle, J., Crook, N., Pal, G., Septer, A. N., Wagner, M. R., Beck, A. E., and Kleiner, M. 2025. Differential metaproteomics of bacteria grown in vitro and in planta reveals functions used during growth on maize roots. 2025.06.02.657423. 10.1101/2025.06.02.657423.

12. Haas, D., and Défago, G. 2005. Biological control of soil-borne pathogens by fluorescent pseudomonads. Nat. Rev. Microbiol. 3:307–319. 10.1038/nrmicro1129.

13. Hemmerle, L., Maier, B. A., Bortfeld-Miller, M., Ryback, B., Gäbelein, C. G., Ackermann, M., and Vorholt, J. A. 2022. Dynamic character displacement among a pair of bacterial phyllosphere commensals in situ. Nat. Commun. 13:2836. 10.1038/s41467-022-30469-3.

14. Jones, D. L. 1998. Organic acids in the rhizosphere – a critical review. Plant Soil 205:25–44. 10.1023/A:1004356007312.

15. Kaya, C., sonmez,Osman, Aydemir,Salih, Ashraf, Muhammad, and and Dikilitas, M. 2013. Exogenous application of mannitol and thiourea regulates plant growth and oxidative stress responses in salt-stressed maize (Zea mays L.). J. Plant Interact. 8:234–241. 10.1080/17429145.2012.725480.

16. Krumbach, J., Kroll, P., Wewer, V., Metzger, S., Ischebeck, T., and Jacoby, R. P. 2021. Metabolic analysis of a bacterial synthetic community from maize roots provides new mechanistic insights into microbiome stability. 2021.11.28.470254. 10.1101/2021.11.28.470254.

17. Li, H., Smith, S. E., Holloway, R. E., Zhu, Y., and Smith, F. A. 2006. Arbuscular mycorrhizal fungi contribute to phosphorus uptake by wheat grown in a phosphorus-fixing soil even in the absence of positive growth responses. New Phytol. 172:536–543. 10.1111/j.1469-8137.2006.01846.x.

18. Lidbury, I. D. E. A., Raguideau, S., Borsetto, C., Murphy, A. R. J., Bottrill, A., Liu, S., Stark, R., Fraser, T., Goodall, A., Jones, A., Bending, G. D., Tibbet, M., Hammond, J. P., Quince, C., Scanlan, D. J., Pandhal, J., and Wellington, E. M.H. 2022. Stimulation of Distinct Rhizosphere Bacteria Drives Phosphorus and Nitrogen Mineralization in Oilseed Rape under Field Conditions. mSystems 7:e00025–22. 10.1128/msystems.00025-22.

19. Marín, O., González, B., and Poupin, M. J. 2021. From Microbial Dynamics to Functionality in the Rhizosphere: A Systematic Review of the Opportunities With Synthetic Microbial Communities. Front. Plant Sci. 12. 10.3389/fpls.2021.650609.

20. Mo, Y., Bier, R., Li, X., Daniels, M., Smith, A., Yu, L., and Kan, J. 2024. Agricultural practices influence soil microbiome assembly and interactions at different depths identified by machine learning. Commun. Biol. 7:1–16. 10.1038/s42003-024-07059-8.

21. Niu, B., and Kolter, R. 2018. Quantification of the Composition Dynamics of a Maize Root-associated Simplified Bacterial Community and Evaluation of Its Biological Control Effect. Bio-Protoc. 8:e2885. 10.21769/BioProtoc.2885.

22. Niu, B., Paulson, J. N., Zheng, X., and Kolter, R. 2017. Simplified and representative bacterial community of maize roots. Proc. Natl. Acad. Sci. 114:E2450–E2459. 10.1073/pnas.1616148114.

23. Niu, X., Chen, M., Tao, A., and Qi, J. 2016. Salinity and Drought Treatment Assays in Kenaf (Hibiscus cannabinus L.). BIO-Protoc. 6. 10.21769/BioProtoc.1918.

24. Parnell, J. J., Vintila, S., Tang, C., Wagner, M. R., and Kleiner, M. 2023. Evaluation of ready-to-use freezer stocks of a synthetic microbial community for maize root colonization. Microbiol. Spectr. 12:e02401–23. 10.1128/spectrum.02401-23.

25. Poudel, M., Mendes, R., Costa, L. A. S., Bueno, C. G., Meng, Y., Folimonova, S. Y., Garrett, K. A., and Martins, S. J. 2021. The Role of Plant-Associated Bacteria, Fungi, and Viruses in Drought Stress Mitigation. Front. Microbiol. 12.

26. Salvato, F., Vintila, S., Finkel, O. M., Dangl, J. L., and Kleiner, M. 2022. Evaluation of Protein Extraction Methods for Metaproteomic Analyses of Root-Associated Microbes. Mol. Plant-Microbe Interactions® 35:977–988. 10.1094/MPMI-05-22-0116-TA.

27. Schäfer, M., Pacheco, A. R., Künzler, R., Bortfeld-Miller, M., Field, C. M., Vayena, E., Hatzimanikatis, V., and Vorholt, J. A. 2023. Metabolic interaction models recapitulate leaf microbiota ecology. Science 381:eadf5121. 10.1126/science.adf5121.

28. van Schaik, J., Li, Z., Cheadle, J., and Crook, N. 2023. Engineering the Maize Root Microbiome: A Rapid MoClo Toolkit and Identification of Potential Bacterial Chassis for Studying Plant–Microbe Interactions. ACS Synth. Biol. 12:3030– 3040. 10.1021/acssynbio.3c00371.

29. Vannier, N., Mesny, F., Getzke, F., Chesneau, G., Dethier, L., Ordon, J., Thiergart, T., and Hacquard, S. 2023. Genome-resolved metatranscriptomics reveals conserved root colonization determinants in a synthetic microbiota. Nat. Commun. 14:8274. 10.1038/s41467-023-43688-z.

30. Vorholt, J. A., Vogel, C., Carlström, C. I., and Müller, D. B. 2017. Establishing Causality: Opportunities of Synthetic Communities for Plant Microbiome Research. Cell Host Microbe 22:142–155. 10.1016/j.chom.2017.07.004.

31. Wagner, M. R., Tang, C., Salvato, F., Clouse, K. M., Bartlett, A., Vintila, S., Phillips, L., Sermons, S., Hoffmann, M., Balint-Kurti, P. J., and Kleiner, M. 2021. Microbe-dependent heterosis in maize. Proc. Natl. Acad. Sci. 118:e2021965118. 10.1073/pnas.2021965118.

32. Wipf, H. M.-L., Bùi, T.-N., and Coleman-Derr, D. 2021. Distinguishing Between the Impacts of Heat and Drought Stress on the Root Microbiome of Sorghum bicolor. Phytobiomes J. 5:166–176. 10.1094/PBIOMES-07-20-0052-R.

33. Yang, J., Kloepper, J. W., and Ryu, C.-M. 2009. Rhizosphere bacteria help plants tolerate abiotic stress. Trends Plant Sci. 14:1–4. 10.1016/j.tplants.2008.10.004.

34. Zhalnina, K., Louie, K. B., Hao, Z., Mansoori, N., da Rocha, U. N., Shi, S., Cho, H., Karaoz, U., Loqué, D., Bowen, B. P., Firestone, M. K., Northen, T. R., and Brodie, E. L. 2018. Dynamic root exudate chemistry and microbial substrate preferences drive patterns in rhizosphere microbial community assembly. Nat. Microbiol. 3:470–480. 10.1038/s41564-018-0129-3.

35. Zhao, D., Raja Reddy, K., Kakani, V. G., Read, J. J., and Carter, G. A. 2003. Corn (Zea mays L.) growth, leaf pigment concentration, photosynthesis and leaf hyperspectral reflectance properties as affected by nitrogen supply. Plant Soil 257:205–218. 10.1023/A:1026233732507.

